# Inhibition of TBK1/IKKe mediated RIPK1 phosphorylation sensitizes tumors to immune cell killing

**DOI:** 10.1101/2025.05.20.655064

**Authors:** Anastasia Piskopou, David W. Vredevoogd, Xiangjun Kong, Daniel S. Peeper, Maarten Altelaar, Kelly E. Stecker

## Abstract

Resistance to immune cell-mediated cytotoxicity poses a significant challenge in cancer therapy, compromising the efficacy of immunotherapeutic approaches such as immune checkpoint blockade (ICB) treatment. To enhance therapy outcomes, it is crucial to identify interventions that can synergize with ICB therapy to overcome tumor resistance. Therefore, we need to define the cellular mechanisms that sensitize tumors to cytotoxic T cells. CD8 T cells rely on cytokines such as TNF to carry out their cytotoxicity against tumors, and recent findings link select tumor mutations in the TNF pathway to increased T cell killing, in a manner dependent on RIPK1 kinase. Here, we demonstrate that sensitized tumor cells fail to initiate inhibitory RIPK1 phosphorylation at site S25 upon T cell attack, thereby foregoing a pro-survival checkpoint early in TNF signal transduction. Consequently, tumor cells experiencing a loss of TNF-induced RIPK1 S25 phosphorylation exhibit increased RIPK1 activation and fail to recruit non-canonical IKK kinases (TBK1 and IKKe) to the TNFR1 complex. Functional knockouts of TBK1 and IKKe in melanoma cells result in heightened sensitivity not only in CD8 T cell but also in Natural Killer cell attacks. Our findings indicate that preventing TBK1 and IKKe recruitment to the TNF signaling complex, thereby blocking RIPK1 pro-survival phosphorylation and promoting direct RIPK1 activation, is a tractable strategy to increase tumor sensitivity to immune cell killing and has the potential to benefit current immunotherapy interventions.

## INTRODUCTION

The implementation of immune checkpoint blockade (ICB) therapy has significantly transformed cancer treatment by focusing clinical interventions on reactivating the body’s immune system to recognize and eliminate cancer cells [1–3]. Drugs such as PDL-1 inhibitors stimulate cytotoxic CD8 T cells, enabling them to reengage with and destroy tumors by removing immunosuppressive signals in the tumor microenvironment [3–5]. Despite promising advancements, most patients experience little durable benefit from ICB therapy due to in part challenges associated with tumor resistance [1, 2, 6, 7]. This highlights the critical need for additional therapy interventions that work alongside immune system activation to combat tumor resistance and restore ICB therapy efficacy. Understanding the mechanisms of intrinsic tumor resistance to immune cell clearance is therefore essential for improving ICB treatment outcomes.

Engaged CD8 T cells release a barrage of cytokines during their attack on target tumor cells, including interferon-gamma (IFNg), tumor necrosis factor (TNF), and TNF-related apoptosis-inducing ligand (TRAIL) [5, 8–10]. For patients responding to ICB therapy, T cell-derived TNF is particularly crucial for mediating cytotoxicity, and increasing tumor sensitivity to TNF can potentially overcome tumor resistance [10]. RIPK1, a key regulator of downstream TNF signaling, has been shown to be essential for CD8 T cell-mediated cytotoxicity, as tumor cells can be driven into RIPK1-dependent apoptosis (RDA) under these conditions [10,11,12]. However, the detailed mechanisms governing RDA in immune cell-tumor interactions remain not yet fully understood.

TNF recognition at the cell surface is mediated predominantly by two receptors: TNF receptor 1 (TNFR1) and TNF receptor 2 (TNFR2). TNFR2 signaling is mainly confined to immune cells, such as T lymphocytes and regulatory T cells, whereas TNFR1 is widely expressed in nearly all cell types, including tumor cells [13]. This broad expression turns TNFR1 into a key focus in tumor studies, as its signaling can either promote cell survival or induce cell death depending on factors such as cell type, TNF dosage, and the recruitment of specific proteins to the receptor complex [14]. TNF signaling outcomes are heavily influenced by the activation status of Receptor Interacting Protein Kinase 1 (RIPK1), a key component of the TNFR1 complex (also referred to as Complex I) [15]. RIPK1 acts as a scaffold protein during Complex I assembly and promotes downstream NF-kB pro-survival inflammatory signaling when inactive [16]. Upon activation, RIPK1 can form cell death protein complexes that initiate either apoptosis or necroptosis [15–20]. The activation of RIPK1 is known to be tightly controlled by post-translational modifications (PTMs), including ubiquitination and phosphorylation, by pro-survival checkpoint kinases within or interacting with the TNFR1 complex [21–23]. Manipulating TNF signaling is inherently complex, as it can lead to both tumor-promoting and tumor-suppressive effects. Both activation and depletion of RIPK1 have been investigated as potential therapeutic strategies in cancer treatment [16, 24–27]. Further research is required to delineate the specific alterations within the TNF signaling pathway that enhance tumor sensitivity to T cell-mediated killing, as well as the precise mechanisms involved.

Herein, we present compelling evidence that TNF-driven T cell-mediated killing relies on direct RIPK1 activation within the TNFR1 complex through the loss of pro-survival RIPK1 phosphorylation. We observed that sensitized tumor cells fail to recruit sufficient levels of pro-survival checkpoint kinases TBK1 and IKKe to Complex I, resulting in the loss of inhibitory Ser25 phosphorylation and an increase in activating Ser166 phosphorylation on RIPK1. Consequently, the direct loss of TBK1 and IKKe in tumor cells leads to heightened sensitivity to both CD8 T cell and natural killer (NK) cell killing. Utilizing four distinct approaches to sensitization that result in TBK1 depletion in the TNFR1 complex, we demonstrate that modulating RIPK1 checkpoint inhibition offers a tractable strategy to unleash TNF toxicity. Our findings suggest that targeting proteins involved in RIPK1 pro-survival checkpoints and promoting direct RIPK1 activation is a viable mechanism to potentiate TNF-induced cell death and enhance the impact of immune cell-mediated tumor clearance.

## RESULTS

### Sensitized tumor cells rapidly activate cell death through TNF signaling and fail to initiate pro-survival phosphorylation events upon T cell challenge

Previous research has demonstrated that TNF alone is not likely to function as a potent antitumor agent under baseline conditions. However, when specific TNF pro-survival signaling components, such as TRAF2, are eliminated, tumors become highly sensitized to TNF’s cytotoxic effects. [10] TRAF2 (TNF Receptor-Associated Factor 2) is a key adaptor protein within the TNFR1 signaling pathway that plays a pivotal role in dictating whether cells undergo survival or cell death in response to TNF stimulation. Within TNFR1 signaling, TRAF2 primarily functions to promote pro-survival pathways by facilitating the recruitment of cellular inhibitors of apoptosis proteins (cIAPs), such as cIAP1 and cIAP2. cIAPs ubiquitinate components of the signaling complex, such as RIPK1, leading to the activation of downstream pro-survival and inflammatory pathways, including NF-κB and MAPK signaling [28,29]. When TRAF2 is depleted, the recruitment of cIAPs to the TNFR1 complex is disrupted, preventing the ubiquitination of RIPK1. In the absence of this modification, RIPK1 transforms into a death-inducing kinase, which can initiate apoptotic or necroptotic cell death pathways [28].

To further explore the molecular mechanisms driving tumor sensitivity to CD8 T cell-mediated killing, we challenged wild-type (WT) and TRAF2 knockout (KO) melanoma cells with healthy donor CD8 T cells using a matched co-culture system. In this system, cells were engineered to express cognate antigen-receptor pairs to enable recognition [10, 30], allowing us to examine the early proteome and phosphoproteome responses.

We first compared within the responses of tumor cells in CD8 T cell (herein referred to as T cell) co-culture to those in direct TNF stimulation. This comparison aimed to determine the extent to which these treatments share a molecular response, given that TNF is proposed to be the primary cytokine responsible for T cell-mediated killing in tumors experiencing heightened T cell challenge [10]. We discovered a striking similarity between the TNF and T cell treatments, with nearly 90% of significantly upregulated phosphorylation sites shared between the two conditions in sensitized TRAF2 KO tumor cells (Fig 1A and 1B). The most substantial phosphorylation induction occurred in pathways controlling cell cycle arrest and ATM-mediated DNA damage response (Fig. 1A and 1C), suggesting rapid activation of extrinsic apoptotic pathways in sensitized tumors [31–34].

**Figure 1.**
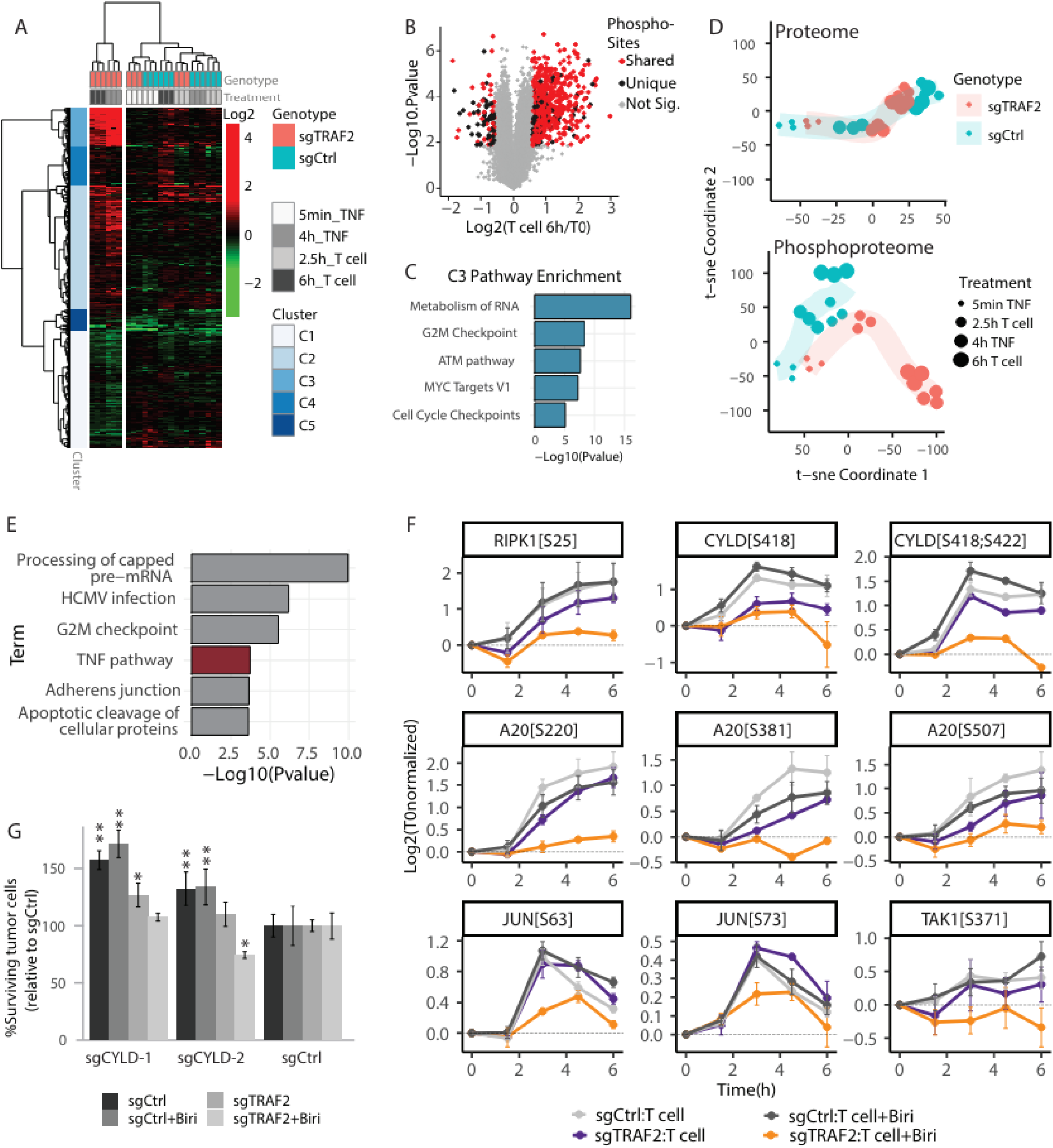
Melanoma cells sensitized to T-cell killing show a distinct phosphorylation response with reduced TNFR1 complex phosphorylation. **(A)** Phosphoproteome clustering of significantly regulated sites following TNF and T cell treatment of polyclonal pools of sgCtrl or sgTRAF2-transduced D10 cells. Values represent T0 normalized phosphorylation changes. **(B)** Overlap of T cell and TNF response in sensitized tumor cells. Phosphorylation changes in sgTRAF2 D10 cells following 6 hours of T cell co-culture vs. sgTRAF2 at T0. Red color coding indicates shared phosphorylation sites that are coordinately regulated in sgTRAF2 D10 cells following 4-hour TNF treatment. **(C)** Enrichment analysis of phosphorylation sites from the most strongly induced phosphorylation cluster in (A). **(D)** t-SNE analysis of significantly regulated proteins and phosphorylation sites in T cell and TNF treatment time course described in (A). **(E)** Enrichment analysis of phosphorylation sites significantly upregulated in sgCtrl BLM cells treated with T cells + Birinipant but not upregulated in sgTRAF2 BLM cells treated under the same conditions. **(F)** Phosphorylation changes in TNFR1-associated proteins from BLM cells treated with T cells +/- Birinipant (1ug/mL). Selected sites show significant upregulation in non-sensitized tumor cells but not in sensitized cells (sensitized cells = orange line; sgTRAF2 + Birinipant). **(G)** CYLD knockout confers resistance to T cell-mediated killing in a WT background. BLM cells were transduced with indicated sgRNAs and subjected to a T cell cytotoxicity assay +/- Birinapant (1ug/mL). Viability was normalized to each condition’s untreated control and then further normalized to each treatment’s sgCtrl sample. Error bars represent standard deviation of biological replicates (n=3). Statistical analysis was performed with a two-way ANOVA with a Dunnet post-hoc test, using genotype and treatment as factors.

Supporting these observations, we found that several kinases involved in DNA damage response, ATM, ATR, and PKCA, showed upregulated kinase activity in sensitized tumor cells following both T cell and TNF treatment (Fig S1). The nearly identical short-term molecular response between TNF and T cell treatment aligns with previous assertions that T cells utilize TNF to execute their cytotoxicity toward tumors. This similarity suggests that both T cell and TNF treatments initiate similar cell death pathways, indicating that the two stimuli may converge on related signaling mechanisms to drive cytotoxicity. Notably, the distinguishing response in sensitized tumors was only observed in the phosphoproteome and not at the proteome level (Fig 1D). This finding suggests that cytotoxic events are initiated before the onset of systematic transcriptional or translational changes, implying that tumors may enact resistance to T cell challenges primarily through modifications at the post-translational level (PTMs) alone.

To identify early phosphorylation events that protect tumor cells from T cell killing we expanded the temporal resolution of our co-culture experiment and examined a more robust mode of tumor sensitization. We modeled this robust sensitization by combining TRAF2 knockout (KO) with drug treatment targeting the TRAF2-interacting proteins cIAP1/2. Dual targeting of TRAF2 and cIAP1/2 has been shown to sensitize a wide range of tumor cell lines across multiple cancer types to T cell killing [10]. Using this information, we performed a matched co-culture study using a melanoma cell line that is only sensitized to T cell killing upon TRAF2 KO and treatment with Birinipant, a SMAC mimetic targeting cIAP1/2 [35]. We analyzed the proteome and phosphoproteome response over five time points spanning six hours. Our hypothesis was that tumor cells in a sensitized state would fail to trigger a defensive response to T cell challenge, resulting in a loss of activation in essential pathways. We specifically examined our data for treatment-induced phosphorylation changes that were upregulated in resistant WT cells over time but were either non-responsive or significantly less responsive in sensitized tumor cells. This analysis identified TNF signaling as significantly affected in sensitized tumors (Fig 1E) and highlighted a loss of treatment-induced phosphorylation in several proteins directly involved in TNFR1 signaling including RIPK1, A20, TAK1, and CYLD (Fig 1F). Except for A20, these proteins exhibited no change in abundance following T-cell treatment (Fig S1B), underscoring the significance of PTM regulation in tumor resistance. RIPK1 was phosphorylated at site S25 in resistant but not sensitized tumor cells in response to T cell challenge (Fig1F). Intriguingly, S25 phosphorylation is described as a pro-survival checkpoint in TNF signaling as it has been shown to confer resistance to RIPK1-mediated cell death by preventing RIPK1 auto-activation in murine models of infection and inflammation [36]. Our data demonstrate that tumor cells are enabling this pro-survival response under T cell challenge, whereas sensitized tumor cells fail to do so. The loss of a RIPK1 pro-survival checkpoint in sensitized tumor cells is particularly informative given that these cells undergo T cell-induced, RIPK1-dependent apoptotic cell death [10]. This directly links the loss of S25 phosphorylation with increased RIPK1 activity and heightened susceptibility to T cell killing.

In addition to RIPK1, we observed increased phosphorylation of CYLD at S418 and S422 in resistant cells (Fig 1F). CYLD is a deubiquitinase (DUB) enzyme that negatively regulates NF-kB signaling and can promote cell death by removing regulatory ubiquitin chains from several TNFR1 complex proteins, including RIPK1 [37–39]. CYLD DUB activity is inhibited by S418 and S422 phosphorylation [40] and thus our data indicates that CYLD inhibition promotes tumor resistance to T cell killing. To validate that loss of CYLD activity results in enhanced resistance to T cells we generated CYLD knockout melanoma lines and tested them in our matched co-culture assay. As predicted, CYLD knockout tumor cells displayed up to 173% improved survival over WT controls when challenged by cytotoxic T cells (Fig 1G). The survival benefit of CYLD depletion appears partially dependent on TRAF2, as TRAF2 KO cells showed minimal to no improved resistance to T cell killing compared to controls (Fig 1G). Overall, these data support that early phosphorylation of TNFR1 signaling complex components is crucial for conferring survival of tumors under T cell challenge and indicate that manipulating RIPK1 checkpoints may be a viable strategy to sensitize tumors to T cell killing.

### Sensitized tumor cells have altered TNFR1 complexes that are deficient in non-canonical IKK kinases

We hypothesized that the loss of T cell-induced phosphorylation of TNF signaling proteins in sensitized tumor cells might stem from changes in the assembly of the TNFR1 signaling complex upon TNF stimulation. To investigate this, we conducted affinity purification of the TNFR1 complex in both sensitized and resistant tumor cells upon short-term stimulation of biotin-labeled TNF and analyzed the samples by LC-MS (Fig 2A). Complex I-associated proteins were significantly enriched above background in our MS data compared to untagged TNF pull-downs used as a negative control, indicating high-quality enrichment of Complex I across our experimental conditions (Fig S2A, S2B). Sensitized tumors harboring both TRAF2 and cIAP1/2 depletions showed significantly reduced recruitment of certain TNFR1 subcomplexes including linear ubiquitin chain assembly complex (LUBAC) proteins, as well as canonical and non-canonical IKK subcomplexes (Fig 2B). The assembly of these components into the TNFR1 complex is known to occur downstream of TRAF2 and cIAP1/2 recruitment and ubiquitination, indicating our findings support current models of TNF signal transduction events [12,41]. Importantly, the kinases hypothesized to phosphorylate RIPK1 at position S25 as part of a pro-survival checkpoint in TNF signaling (IKKa, IKKb, TBK1 and IKKe) were all significantly reduced in the TNFR1 complex of sensitized tumors. Deceased levels of these kinases align with our observations of decreased RIPK1 S25 phosphorylation following T cell treatment in sensitized tumor cells, indicating that loss of checkpoint kinase recruitment to complex I may be a putative mechanism driving tumor sensitivity to T cell attack.

**Figure 2.**
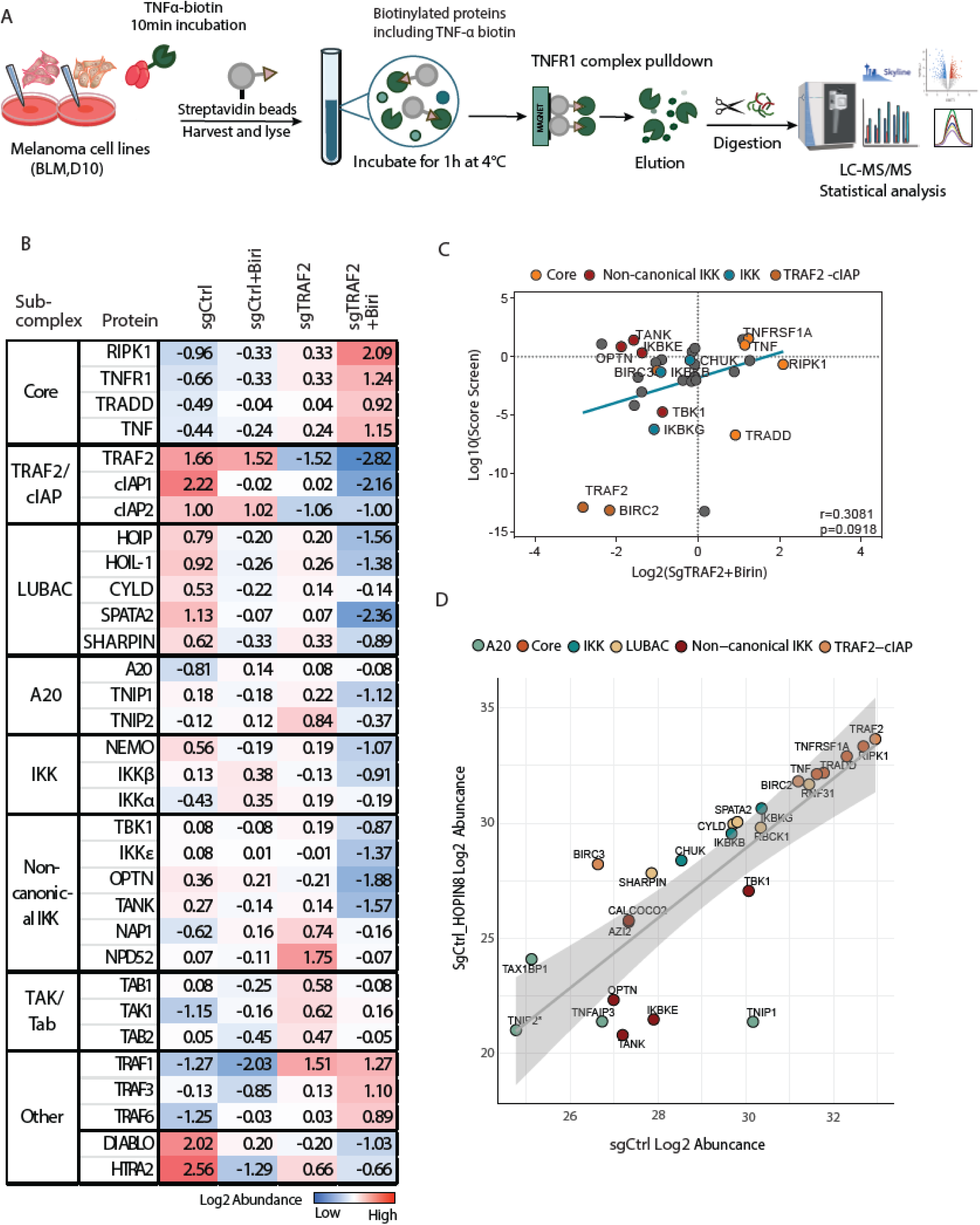
Sensitized tumor cells display altered TNFR1 complexes deficient in pro-survival IKK checkpoint kinases. **(A)** Overview of the experimental workflow. **(B)** Relative abundance of Complex I proteins measured using TNF-biotin affinity purification and mass spectrometry (AP-MS) under the 4 listed treatment conditions. Values are Log2 transformed, median normalized, and represent the average of 3 biological replicates per treatment and with missing values imputed. **(C)** Correlation between relative protein abundance in Complex I in sensitized tumor cells and survival score in T cell sensitivity screening. Reduced protein abundance in Complex I correlates with increased sensitivity to T cell-mediated killing in sgRNA drop-out screens [10]. Negative ratios (X-axis) indicate decreased abundance in the TNFR1 complex in sensitized tumor cells (sgTRAF2 + Birinipant) relative to WT controls and negative screen hits (Y-axis) correspond to lower tumor cell survival upon T cell challenge. **(D)** Correlation between relative protein abundance of Complex I proteins in sensitive vs resistant tumor cells. Tumor cells were sensitized to T cell-mediated killing through treatment with RNF31 inhibitor HOPIN8 and Complex I proteins were measured by TNF-biotin AP-MS. LFQ intensity values are Log2 transformed and represent the average of 3 biological replicates per condition with missing values imputed. Complex I proteins are color-coded by subcomplex. Panel **A** was partially created with <u>BioRender.com</u> and <u>bioicons.com</u>.

To determine if the altered recruitment of proteins to the TNFR1 complex is a broader mechanism through which tumors can be sensitized to T cell killing, we compared the relative protein abundance of TNFR1 complex proteins in sensitized vs. WT tumor cells with enrichment scores from sgRNA knockout screens that identify proteins whose genetic ablation increases sensitivity to T cell killing [10]. Our analysis revealed a positive correlation between the depletion of proteins in the TNFR1 complex and their capacity to sensitize tumor cells to T-cell attack upon knockout (Fig 2C). Specifically, proteins that showed the most significant reduction in recruitment to Complex I also had the greatest negative impact on cell viability following T cell challenge when genetically disrupted. This suggests that tumors can be driven into a ‘sensitized state’ by disrupting the TNFR1 complex through various approaches that result in TNF signaling outcomes which favor cell death in response to T cell challenge.

To further elucidate the essential recruitment events within the TNFR1 complex that prevent T cell-mediated killing in tumor cells, we examined the TNFR1 complex composition in melanoma cells with a knockout mutation in RNF31 [42]. RNF31 functions downstream of TRAF2 and cIAP1/2 within the TNFR1 complex as part of the LUBAC subcomplex, which adds M1-linked linear ubiquitin chains to proteins, including RIPK1. These ubiquitin chains stabilize the complex and facilitate the recruitment of additional proteins, such as IKK checkpoint kinases. Our data from TRAF2 KO combined with Birinipant treatment indicated a significant depletion of RNF31 in the TNFR1 complex in sensitized tumor cells, despite unchanged cellular RNF31 protein levels (Fig 2B, Fig S2E-F). Genetic screens further showed that knocking out RNF31 increases sensitivity to T cell killing (Fig 2C, [10, 42]. RNF31 ubiquitination activity was shown to be selectively inhibited by the drug HOPIN-8 [43] and drug treatment which sensitizes tumors to T cell killing in vivo to the same extent as RNF31 KOs [42].

To understand the mechanism behind tumor sensitization we compared the TNFR1 complex in WT and RNF31 KO tumor cells with and without HOPIN-8 drug treatment. Our analysis in previous studies revealed that RNF31 KOs and HOPIN-8 treatment significantly reduced the recruitment of A20 and non-canonical IKK subcomplexes to the receptor (Fig 2D, S2C, S2D). A20 plays a critical role in removing K63-linked polyubiquitin from RIPK1 and other components of the TNFR1 signaling complex, thereby regulating signal transduction and influencing the balance between cell survival and death [44]. The loss of A20 recruitment in both conditions suggests that ubiquitination dynamics within the TNFR1 complex are disrupted, tipping the balance toward pro-death signaling. While RNF31 KOs led to the depletion of many more proteins in the TNFR1 complex, the reduction of A20 and non-canonical IKK subcomplexes was the only consistent change across both sensitizing conditions (Fig 2D, S2C, S2D). Moreover, in sensitized tumor states caused by TRAF2 and cIAP1/2 depletion, the loss of non-canonical IKK proteins in the TNFR1 complex emerged as the sole shared feature. Together this data indicates that preserving recruitment of TBK1 and IKKe kinases along with their associated scaffolding proteins to the TNFR1 complex, is essential for tumor cells to resist T cell attack.

### Sensitized tumor cells fail to inhibit RIPK1 activation in the TNFR1 complex

To determine whether the phosphorylation state of RIPK1 within the TNFR1 complex correlates with the recruitment of checkpoint kinases, we examined RIPK1 phosphorylation levels in our TNFR1 purifications, using AP-MS. We analyzed the TNFR1 complex dynamics across 8 conditions, representing 4 sensitized and 4 resistant tumor states. We observed consistently high levels of RIPK1 S25 phosphorylation in WT cells and a loss of S25 phosphorylation in sensitized cells (Fig 3A and 3B). These findings align with our T cell co-culture time course data (Fig 1F). Moreover, sensitized tumor cells exhibited a corresponding increase in RIPK1 S166 phosphorylation within the TNFR1 complex. Phosphorylation at S166 is indicative of RIPK1 autoactivation and is shown to be required for RIPK1-mediated cell death in several mouse models of inflammatory disease [45]. To validate our findings, we developed a targeted MS method (SRM) that allows us to measure RIPK1 protein abundance and phosphorylation at several positions. We confirmed that S25 and S166 phosphorylation exist in opposing occupancy in the TNFR1 complex of sensitized and resistant cells, while phosphorylation at a third site, S320, remains constant (Fig 3C).

**Figure 3.**
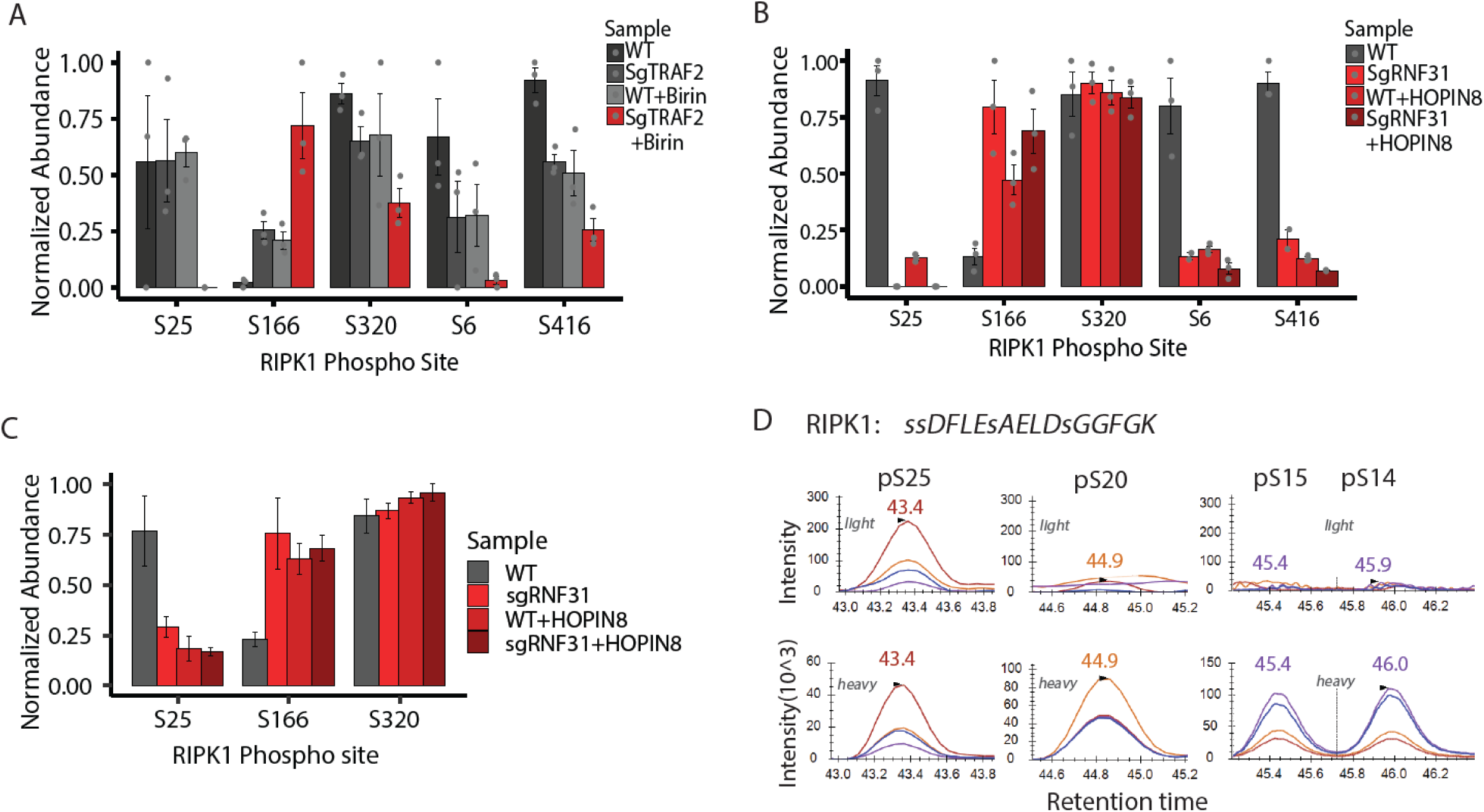
RIPK1 S25 phosphorylation in the TNFR1 complex is deficient in tumor cells that are -sensitive to T cell-mediated killing. **(A)** Phosphorylation of RIPK1 within Complex I measured following TNF-biotin AP-MS. Bar color codes indicate tumor sensitivity to T cell killing (grey scale= resistant; red scale = sensitized). Error bars indicate biological replicate standard deviation (n=3). Phosphopeptide abundances were first normalized to endogenous RIPK1 protein levels within Complex I and then normalized to the maximum value for each site across experimental conditions. **(B)** Same as in (A) but tumor cells were sensitized to T cell-mediated killing through knocking out RNF31 and/or treatment with RNF31 inhibitor HOPIN8. **(C)** Same as in (B) but RIPK1 peptides were measured by SRM using heavy-labeled synthetic peptide standards for quantification. Peptide sequences and transitions used for quantification are listed in Table S1. **(D)** SRM extracted ion chromatograms for all RIPK1 phospho-isoforms of the tryptic peptide containing S25. Colored traces represent different transitions for each phospho-isoform, the upper panels show endogenous signal from a WT sgCtrl sample, the lower panels show the corresponding intensity from heavy labeled synthetic peptide standards.

Interestingly, S25 phosphorylation remains low in WT + HOPIN-8 treated samples, where IKKa and IKKb kinases are present at WT levels, but TBK1 and IKKe kinases are significantly reduced in the TNFR1 complex. This suggests that the non-canonical IKKs, TBK1 and IKKe, are the checkpoint kinases responsible for RIPK1 S25 phosphorylation. Given that S25 phosphorylation occurs on a peptide sequence containing three additional annotated phosphorylation sites (Figure 3D), we confirmed the localization of S25 using synthetic standards for adjacent S20, S14, and S15 phosphorylation, finding that the endogenous signal was detected only at site S25 (Fig 3D). Overall, our RIPK1 phosphorylation results reveal a pattern in which sensitized tumor cells lose pro-survival S25 phosphorylation and gain cell-death-promoting S166 phosphorylation under conditions where non-canonical IKKs fail to be recruited to the TNFR1 complex upon TNF stimulation.

### Loss of non-canonical IKK checkpoint kinases sensitizes tumors to immune cell killing

TBK1 and IKKe kinases are recruited to the TNFR1 complex downstream of TRAF2, cIAP1/2 and RNF31 and their presence in the complex appears to be essential for tumors to resist T cell challenge [46]. Therefore, we aimed to investigate whether knocking out these kinases directly enhances tumor susceptibility to T cell killing. We generated polyclonal functional TBK1 KOs in two different melanoma cell lines (BLM, D10) and observed that loss of TBK1 alone is able to sensitize cells to T cell challenge in some melanoma cell lines (Fig 4A and 4B), while knocking out both TBK1 and IKKe is required in others (Fig 4C). Significantly, TBK1 and TBK1+IKKe KOs showed heighten sensitivity not only to T cells, but also NK cells, suggesting their involvement in a broader mechanism of tumor susceptibility to immune cell attack (Fig 4B and 4C). To confirm that the observed cell death operates through the TNF pathway, as indicated by our previous findings, we conducted parallel tests using TNF treatment and found that TBK1 KOs were sensitized to TNF but not to IFNgamma (Fig 4A). Finally, to contextualize TBK1/IKKe KOs with previously described sensitizing mutations affecting the same signaling pathway, we compared our samples to RNF31 KOs and observed that TBK1 and IKKe depletion sensitized tumor cells to T cell attack to an even greater extent than RNF31 depletion with these T cell donors (Fig 4C).

**Figure 4.**
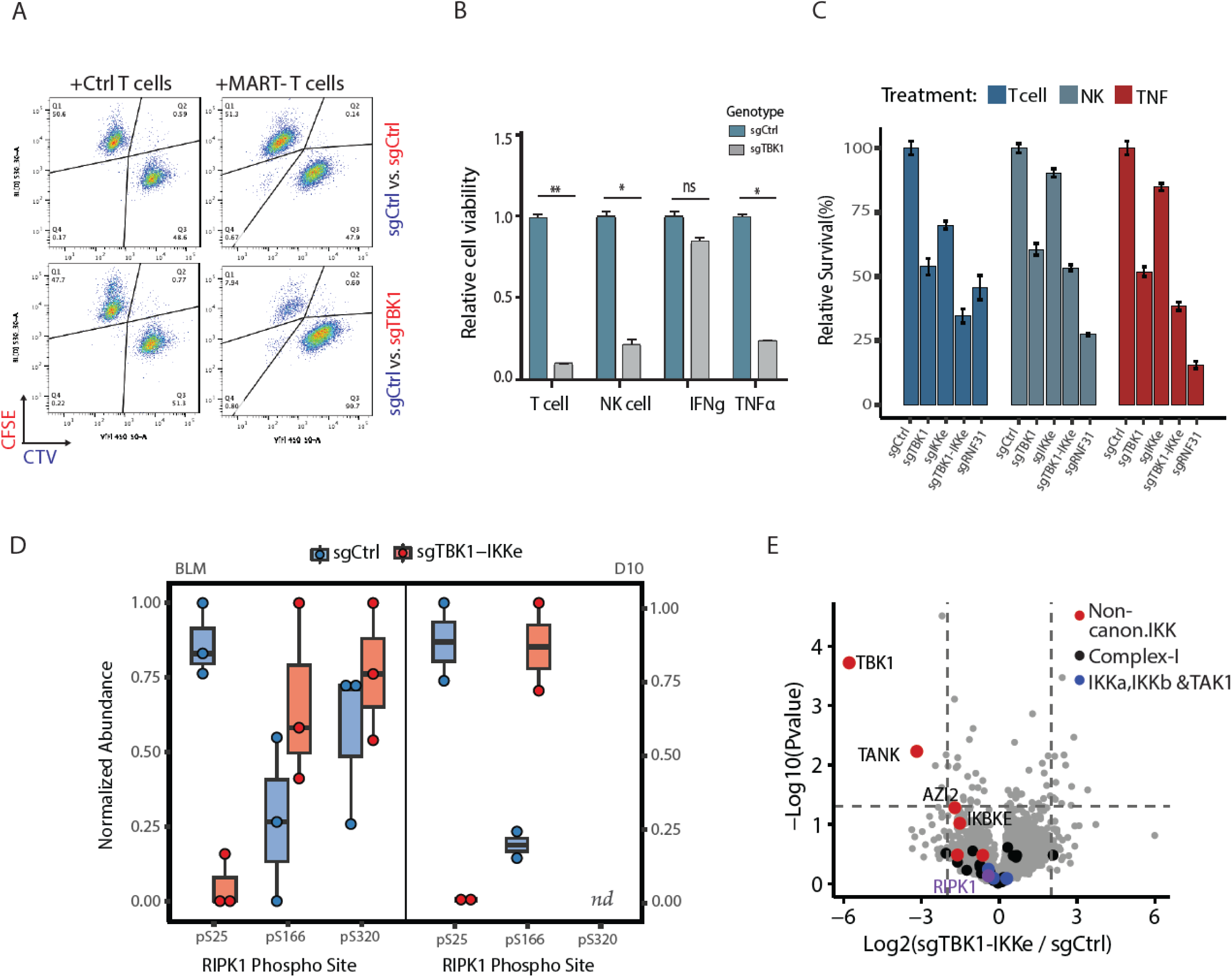
Loss of non-canonical IKKs sensitize tumor cells to immune cell killing. **(A)** Flow cytometry plot of the competition assay of sgCtrl vs. sgTBK1 transduced D10 cells co-cultured with either MART-1 or Ctrl T cells for 3 days. **(B)** Quantification of the competition assay in (A) and additional conditions listed. Treatments were applied as follows: 1:8 T cell: tumor cell ratio; 1:2 NK cell: tumor cell ratio; 5ng/ml INFgamma; 50ng/ml TNF. Cells were sorted after 3 days. Error bars represent standard deviation between biological replicates (n=3). **(C)** Competition assay performed the same as in (A) and (B) but using BLM cells lines transduced with sgCtrl, sgTBK1, sgIKKe, sgTBK1+sgIKKe, or sgRNF31. Error bars represent standard deviation between biological replicates (n=3). **(D)** RIPK1 phosphorylation in the TNFR1 complex measured in sgCtrl and sgTBK1+sgIKKe melanoma cell lines following TNF-biotin AP-MS. Dots represent biological replicates (n=3 for BLM, n=2 for D10). RIPK1 S320 phosphorylation was not detected in D10 cells. Phosphopeptide abundances were first normalized to endogenous RIPK1 protein levels within Complex I and then normalized to the maximum value for each site across experimental conditions. **(E)** Relative abundance of Complex I proteins in sgCtrl vs sgTBK1+sgIKKe tumor cells following TNF-biotin AP-MS. Horizontal line indicates a P-value < 0.05, vertical lines indicate fold-change > 4, n=3 biological replicates per condition.

Our accumulated evidence suggests that the loss of TBK1 and IKKe may sensitize tumor cells to T cell killing by impairing the activation of pro-survival RIPK1 S25 phosphorylation within the TNFR1 complex, thereby promoting RIPK1 activation and subsequent cell death. To confirm that TBK1/IKKe KOs display a corresponding loss of RIPK1 S25 phosphorylation upon TNF stimulation we performed purification of the TNFR1 complex following TNF treatment and observed that RIPK1 S25 phosphorylation was nearly absent in TBK1/IKKe knockouts (Fig 4D). Additionally, TBK1/IKKe knockouts of both melanoma cell lines exhibited increased RIPK1 S166 phosphorylation, indicating heightened RIPK1 activation within the TNFR1 complex compared to WT controls. Importantly, there was no difference in the recruitment of RIPK1, TAK1, or canonical IKKs (IKKa and IKKb) to the complex between TBK1/IKKe KO and WT controls (Fig 4E). This data suggests that the loss of TBK1 and IKKe noncanonical checkpoint kinases is specifically responsible for the reduction in RIPK1 pro-survival phosphorylation. Given that RIPK1 S25 levels were not completely absent in all TBK1/IKKe KOs, it may be that there is a low level of redundancy in phosphorylating this RIPK1 site by other checkpoint kinases. It is also possible that our polyclonal knockouts do not achieve complete depletion of functional TBK1 or IKKe protein levels, despite falling below the limit of detection. Taken together, our data demonstrates that the absence of TBK1 and IKKe sensitizes tumor cells to T cell killing and is directly linked to loss of RIPK1 S25 pro-survival phosphorylation upon TNF stimulation.

## DISCUSSION

Overcoming tumor resistance to immune cell-mediated cytotoxicity remains a critical challenge in improving the efficacy of cancer immunotherapies such as immune checkpoint blockade (ICB) therapy. Central to this resistance is the TNF signaling pathway, which acts as a critical switch between cell survival and cell death. Our findings underscore the pivotal role of RIPK1 within the TNFR1 complex as a key regulator of this balance, heavily influenced by post-translational modifications (PTMs) such as phosphorylation and ubiquitination [10,19,46]. Among the regulatory mechanisms of RIPK1, we identified two non-canonical kinases, TBK1 and IKKe, that serve as crucial checkpoints, mediating pro-survival phosphorylation of RIPK1. Inhibiting TBK1/IKKe prevents RIPK1 S25 phosphorylation, shifts TNF signaling toward cell death, and sensitizes tumors to CD8+ T cell attacks.

Our AP-MS analysis revealed diminished recruitment of ubiquitin-related machinery, including LUBAC proteins, in sensitized tumor cells. Ubiquitination is essential for stabilizing TNFR1 complex assembly and facilitating downstream signaling. Yet, our data suggest that ubiquitin modifications are not the only regulatory layer, as non-canonical ubiquitination likely contributes to RIPK1 phosphorylation and the recruitment of key kinases, underscoring the multifaceted regulatory mechanisms controlling TNF signaling [47]. The activity of CYLD deubiquitinase, was shown to be inhibited in resistant tumor cells via phosphorylation at S418 and S422. This may suggest that tumor cells can evade immune-mediated killing by suppressing CYLD activity and stabilizing pro-survival signaling within the TNFR1 complex. Genetic ablation of CYLD, significantly enhanced resistance to T cell killing, confirming its role as a critical regulator of immune resistance. However, while we see a clear depletion of ubiquitin machinery components in sensitized states, we cannot conclusively determine the precise ubiquitin linkages or dynamics in the absence of more direct assays.

In-depth investigation of the TNFR1 complex dynamics composition showed that sensitized tumor cells have altered assemblies that are deficient in key pro-survival components. This was evident across various methods of sensitization, including TRAF2 knockout combined with cIAP1/2 inhibition as well as RNF31 knockout as previously reported [42]. The consistent finding across these conditions was the diminished presence of TBK1 and IKKe non-canonical kinases within the TNFR1 complex, which points to their direct influence on the phosphorylation status of RIPK1. While canonical kinases (IKKα/β) have been extensively studied in mouse models, understanding their precise role in human TNFR1 signaling is not completely defined. Canonical IKKs are central to NF-κB signaling, promoting cell survival and inflammation. However, their role appears to be context-dependent, varying by species and stimuli. In murine models, TBK1 and IKKε have been shown to negatively regulate canonical IKK activity, suggesting a complex interplay between these kinases [21]. This raises the intriguing possibility that in specific contexts, particurarly in human tumors, the pro-survival signaling mediated by TNF may rely more heavily on TBK1/IKKe than on canonical kinases, as supported by our findings. By focusing on the role of TBK1 and IKKε in regulating RIPK1-mediated cell death, our study provides mechanistic insight into how tumors resist immune-mediated killing and identifies an actionable vulnerability in the TNFR1 signaling axis. This is further reinforced by studies showing that TBK1 depletion sensitizes tumors to T cell cytotoxicity and enhances the efficacy of immune checkpoint blockade [46,48]. Moreover, a recently published study identifies TBK1 and IKKε as suppressors of inflammation downstream of RIPK1 and NLRP3, highlighting their broader role in restraining immune responses triggered by programmed cell death signaling pathways [49].

When comparing strategies for manipulating RIPK1, direct activation of RIPK1 through the loss of checkpoint kinases such as TBK1/IKKe may exhibit higher efficiency in tumor killing than knockout strategies, such as RIPK1 KO or PROTAC-mediated degradation, which eliminate the protein entirely. Although removing RIPK1 altogether prevents its involvement in pro-survival signaling, it also forfeits any potential to leverage its death-inducing functions. In contrast, promoting RIPK1 activation by removing pro-survival phosphorylation checkpoints (pS25) while enhancing pro-death phosphorylation (pS166) offers a more targeted approach that bypasses tumor survival pathways. By disrupting TBK1/IKKe recruitment to the TNFR1 complex, the balance can be shifted from survival to death, overcoming tumor resistance. Collectively, our findings suggest that disrupting pro-survival phosphorylation checkpoints within TNF signaling emerges as a promising strategy to sensitize tumors to the cytotoxic effect of immune cells, potentially enhancing the efficacy of current immunotherapy approaches.

## LIMITATIONS OF THE STUDY

Our study predominantly employs in vitro models to explore the interactions between TNF and T cells with sensitized tumor cells. While this approach provides valuable insights into early signaling events, it does not fully account for the complexities of the tumor microenvironment in vivo. It is important to note that while we examined phosphoproteomic changes, no direct ubiquitin measurement was performed in this study. Finally, further insights are needed into the activation state of canonical IKKα/β kinases and the potential interplay between canonical and non-canonical pathways in the observed sensitized states.

## ACKNOWLEDGMENTS

This work received support from the Horizon 2020 program INFRAIA project Epic-XS (Project 823839) and the NWO-funded Netherlands Proteomics Center through the National Road Map for Large-scale Infrastructures program XOmics (Project 184.034.019). We would also like to thank the flow cytometry facility in NKI-AVL for contributing to this work. Finally, we want to acknowledge funding support from NWO (OCENW.M.22.233).

## METHODS

### Human primary CD8+ T cells and NK cells

Peripheral blood mononuclear cells (PBMCs) were isolated from fresh buffy coats obtained from healthy donors (Sanquin, Amsterdam, the Netherlands) using Ficoll-Paque PLUS density gradient medium (17144003, GE Healthcare) via density gradient centrifugation. Donor buffy coats (B2825R00) were provided by Sanquin, where the clinical information of the donors was recorded. CD8+ T cells were purified from the PBMC fraction using CD8 Dynabeads (Thermo Fisher Scientific) following the manufacturer’s instructions. Isolated CD8+ T cells were activated for 48 hours in 24-well plates pre-coated overnight with CD3 and CD28 antibodies (eBioscience, 5 μg/well) at a density of 2 × 10⁶ cells/well. Subsequently, 2 × 10⁶ activated CD8+ T cells were spinfected with a 1:1 MART-1 TCR retrovirus mixture on Retronectin-coated (Takara, 25 μg/well) non-tissue-culture-treated 24-well plates for 2 hours at 2000 × g. After a 24-hour infection period, the MART-1 T cells were harvested and cultured for 7 days, with TCR expression confirmed via flow cytometry (BD PharMingen, α-mouse TCR β chain).

CD8+ T cells were initially cultured in RPMI medium (GIBCO) supplemented with 10% human serum (One Lambda), 100 U/mL penicillin, 100 μg/mL streptomycin, 100 U/mL IL-2 (Proleukin, Novartis), 10 ng/mL IL-7 (ImmunoTools), and 10 ng/mL IL-15 (ImmunoTools). Post-retroviral transduction, T cells were maintained in RPMI medium containing 10% fetal bovine serum (FBS) and 100 U/mL IL-2. For natural killer (NK) cell isolation and expansion, PBMCs were co-cultured with gamma-irradiated Jurkat cells in RPMI-1640 medium supplemented with 100 U/mL IL-2 for 7 days. NK cells were then purified using the Dynabeads™ Untouched™ Human NK Cells Kit (11349D, Thermo Fisher Scientific). Isolated NK cells were cultured in RPMI-1640 medium containing 9% FBS and 100 U/mL IL-2. All study procedures were approved by the local Institutional Review Board committee.

### Genome-wide CRISPR-cas9 knockout (GeCKO) screen

Human D10 melanoma cells (600 million) were transduced with GeCKO libraries (A and B) using lentiviral vectors at a multiplicity of infection (MOI) of 0.3 as previously described [42]. Two days post-infection, cells were selected with puromycin (2 μg/mL) for three days, and T0 samples were collected. One week later, the cells were divided into two groups: one group underwent co-culture with MART-1 CD8+ T cells (T cell:tumor ratio of 1:8), while the other was left untreated. After two days of co-culture, cells were washed twice with PBS and cultured for an additional four days. The surviving cells were then harvested, and genomic DNA was extracted using the Blood and Cell Culture MAXI kit (13362, Qiagen). sgRNA sequences were amplified by PCR using NEBNext High-Fidelity 2x PCR Master Mix (M0541L, New England BioLabs) with the following primers:

GeCKO forward:

5′-AATGATACGGCGACCACCGAGATCTACACTCTTTCCCTACACGACGCTCTTCCGATCTNNNNNNGGCTTTATATA TCTTGTGGAAAGGACGAAACACC-3′

GeCKO reverse:

5′-CAAGCAGAAGACGGCATACGAGATCCGACTCGGTGCCATTTTTCAA-3′

The six N nucleotides in the forward primer represent unique barcodes for sample identification. Amplified PCR products were purified, pooled in equimolar amounts, and subjected to deep sequencing on the Illumina HiSeq 2500 System. Sequence reads were aligned to the GeCKO A and B libraries, and counts per sgRNA were generated. Reads containing mismatches in sgRNA or common sequences were excluded from analysis. Gene enrichment and depletion at the sgRNA level were determined using the MAGeCK algorithm (v0.5.9.4). Enrichment and depletion in CD8+ T cell and NK cell samples were calculated relative to untreated controls. Results were further analyzed with the MAGeCKFlute package (v1.10.0) in R. Genes with MAGeCK RRA scores corresponding to a -log10 transformed RRA score of ≥3 were considered significantly enriched or depleted.

### Cell lines

D10 (ATCC) and BLM (ATCC) cells were cultured in Dulbecco’s Modified Eagle’s Medium (DMEM) supplemented with 10% fetal bovine serum (FBS, Sigma) and 2 mM glutamine. Regular mycoplasma testing was conducted to ensure contamination-free cultures. All cells were maintained in a humidified incubator at 37°C with 5% CO2. Experiments were conducted between the 5th and 10th passages to minimize cell heterogeneity.

### Competition assays

Cells containing sgRNAs of interest were labeled either with CellTrace CFSE (CFSE, C34554, ThermoFisher) or CellTrace Violet (CTV, C34557, ThermoFisher), mixed at 1:1 ratio and plated in 6-well plates (1×106 D10 cells per well) in triplicate. Labeled cells were challenged with either MART-1 CD8+ T cells (E:T of 1:8), primary NK cells (1:2), IFNg (5ng/ml) or TNF (50ng/ml) for 3 days. Cells were then washed with PBS twice analysed by FACS.

### Flow cytometry

Cells were stained with antibodies targeting surface or intracellular molecules of interest according to manufacturer’s instructions and analyzed on a Fortessa flow cytometer (BD Bioscience). Antibodies against mouse TCR β:Hamster-PE (1:200 dilution, Biolegend, H57-597), human CD8:PacificBlue (1:50 dilution, BD, RPA-T8), human CD8:BB515 (1:100 dilution, BD, RPA-T8), human CD4:PerCP-eF710 (1:100 dilution, eBioscience, SK3), human CD25:PE-Dazzle 594 (1:100 dilution, Biolegend, M-A251), human CD127:Brilliant Violet 650 (1:100 dilution, Biolegend, A019D5) and human FoxP3:APC (1:100 dilution, eBioscience, 236A1E7) were used. The dyes Life/Dead (1:1000 dilution, Themo Fisher) and CTV (1:1000 dilution, Themo Fisher) were used to stain cells.

### Immunoblotting

Cell lysates were denatured and reduced in XT Sample buffer (Bio-Rad) supplemented with 25 mM DTT at 95°C for 5 minutes. Proteins were then separated on a 12% Bis-Tris SDS-PAGE gel (Bio-Rad) and electroblotted onto PVDF membranes. The membranes were blocked with 5% non-fat milk in TBS 1X supplemented with 0.1% Tween 20 (TBS-T) and incubated overnight at 4°C with the following primary antibodies: RIPK1 (#3493, CST), TBK1 (#5483, CST), IKKe (#8644, CST), and β-actin (#4970, CST). Subsequently, membranes were incubated with an anti-rabbit horseradish peroxidase conjugate (#7074, CST) as the secondary antibody for 1 hour at room temperature. When multiple primary antibodies were used, membranes were stripped using Restore PLUS Western Blot Stripping Buffer (Pierce) for 7 minutes and re-incubated as previously described. Detection was carried out using enhanced chemiluminescent substrate (Pierce) and measured with the Amersham Imager 600 (GE Healthcare Bio-Sciences).

### SgRNA construction and lentivirus production

For the construction of a sgRNA-expressing vector, DNA oligonucleotides were annealed and ligated into BsmBI-digested LentiCRISPRv2 plasmid. Target sgRNA oligonucleotide sequences are listed in Table S4. Double knockouts were achieved by employing both puromycin-selectable and blasticidin-selectable variants of lentiCRISPR-v2 for each sgRNA.

Briefly, HEKT293T cells were used for lentivirus production containing mutation and SgRNA control constructs, or a GFP-positive control to determine transfection efficiency. Medium-containing pseudo-virus was harvested and concentrated 2- and 3-days post-transfection. Melanoma cells were seeded in a 6-well plate in a seeding density of 2.5 × 104 per well, where different virus volumes ranging from 2.5 to 15 μl were added. Transduction was enhanced using Polybrene® (Sigma-Aldrich) to neutralize charge interactions and increase binding between the pseudoviral capsid and the cellular membrane. A lethal dose of 10 μg/ml of puromycin (Sigma-Aldrich) was added 3 days post-transduction. Medium with puromycin was refreshed every 2 days until puromycin-resistant colonies of melanoma cells appeared.

### TNF co-immunoprecipitation and sample preparation

1×10⁷ BLM or D10 cells (sgCtrl or sgTBK1/IKKe) were cultured for 24 hours and subsequently incubated with 100 ng/mL biotinylated TNF (BT210–010, R&D Systems) or unlabeled TNF (300-01A, PeproTech) for 10 minutes. Following this treatment, cells were harvested and lysed using a homemade IP lysis buffer (30 mM Tris-HCl pH 7.4, 120 mM NaCl, 2 mM EDTA, 2 mM KCl, 1% Triton X-100) containing a protease and phosphatase inhibitor cocktail (Roche). Bradford protein assay (Bio-Rad Protein Assay Kit I, Bio-Rad) was used to quantify the protein content of replicates for normalization purposes. TNF receptor complexes were precipitated by incubating the lysates with streptavidin-coated magnetic beads (#88816, Thermo Fisher Scientific) for 1 h at 4°C. Purified protein complexes were eluted from magnetic beads using a 5% SDS solution and were digested using S-TRAP microfilters (C02-micro-10, ProtiFi) as previously described [50]. Briefly, eluted samples were reduced and alkylated using DTT (20 mM, 10 min, 95°C) and IAA (40 mM, 30 min). The samples were acidified and proteins were precipitated using a methanol TEAB buffer before loading on the S-TRAP column. After washing the trapped proteins four times with the methanol TEAB buffer, digestion was conducted for 1 h 20 min at 47°C using a total of 1 μg <u>Trypsin</u> (Promega) per sample. Digested peptides were eluted and dried in a vacuum centrifuge before LC-MS analysis.

### LC-MS/MS

For mass spectrometry analysis, spectral data were acquired with an Orbitrap Exploris 480 mass spectrometer (Thermo Fisher Scientific, USA) or an Orbitrap HFX mass spectrometer (Thermo Fisher Scientific, USA) coupled to Ultimate3000 (Thermo Fisher Scientific) UHPLC. The peptides were trapped on an Acclaim Pepmap 100 C18 (5 mm × 0.3 mm, 5 μm, ThermoFisher Scientific) trap column and separated on an in-house packed analytical column (75 µm, ReproSil-Pur C18-AQ 2.4 µm resin (Dr. Maisch GmbH). Solvent B consisted of 80% acetonitrile in 0.1% FA. Trapping of peptides was performed for 2 min in 9% B followed by peptide separation in the analytical column using a gradient of 13 to 44% B in 95 min. After peptide separation, gradients were followed by a steep increase to 99% B in 3 min, a 5-min wash in 99% B, and a 10-min re-equilibration at 9% B. Flow rate was kept at 300 nL/min. MS data was acquired using a DDA method at a resolution of 60,000 and a scan range of 375–1600 m/z. Automatic gain control (AGC) target was set to 3E6, under standard calculation of maximum injection time. The cycle time for MS2 fragmentation scans was set to 1 s. Peptides with charged states 2 to 10 were fragmented, with a dynamic exclusion of 16 s. Fragmentation was done using stepped higher-energy collisional dissociation (HCD)-normalized collision energy of 28%. Fragment ions were accumulated until a target value of 1 × 105 ions was reached or under a maximum injection time of 300 ms, with an isolation window of 1.4 m/z before injection in the Orbitrap for MS2 analysis at a resolution of 15,000. Analysis of melanoma co-cultures with donor-derived T-cells (Figure 1 pannel) was conducted as previously described [10].

### Phosphopeptide Enrichment

Phosphopeptides were enriched in an automated fashion using the AssayMAP Bravo Platform (Agilent Technologies) with Fe(III)-NTA 5 μl cartridges (Agilent Technologies), as previously described [50]. Samples were dissolved in 200 μL of loading buffer (80% acetonitrile/0.1% TFA). Cartridges were primed with 100 μl of priming buffer and equilibrated with 250 μl equilibration buffer (80% ACN, 0.1% TFA). A total of 200 μl per replicate was loaded onto the cartridges and washed twice with equilibration buffer. Elution of phosphopeptides was performed with 2.5% ammonia solution, into 35 μL of 10% formic acid. Eluates were subsequently vacuum-dried and stored at -80 °C until SRM or shotgun LC-MS analysis.

### SRM Assay Development

The development of the SRM assay was guided by an in-house generated spectral library, confirming the synthesized peptide sequences and providing essential LC and MS characteristics for each peptide. This library facilitated the assay development for non-modified and phosphorylated RIPK1 peptides (JPT Peptide Technologies). Using Skyline (v20.2), the 10 most abundant transitions found in the spectral library were selected for the initial method development. The concentration of each heavy isotope-labeled peptide standard was optimized to achieve Gaussian distribution of MS peaks. From these initial measurements, the 5 transitions with the highest area under the curve (AUC) were chosen for further method development. Collision energies (CE) were optimized for each peptide fragment, based on instrument-specific parameters predicted by Skyline (CE = 0.034m/z + 3.314 for doubly charged ions, CE = 0.044m/z + 3.314 for triply charged ions) [51]. Each peptide fragment underwent 11 scans on a TSQ Altis Triple Quadrupole Mass Spectrometer (Thermo Scientific™), with CE values ranging ±5 V around the predicted value. The three most abundant transitions for each peptide were then selected, and a retention window of 4 minutes was established to enable high dwell times at low cycle times (2 seconds). Spectra from SRM MS measurements were analyzed using Skyline as previously described [52]. Signal quality was assessed based on the similarity of sequence and chemical properties between endogenous and heavy-labeled peptides. Inclusion criteria included perfect co-elution, peak shape, similarity of relative transition intensities, and an rdotp ≥ 0.9.

### Data-dependent analysis

RAW files were processed with MaxQuant (version 1.6.10), or Proteome Discoverer (version 2.3.0) MS/MS spectra were searched against the UniProt human database (9606, Homo Sapiens) including common contaminants. Trypsin digestion and a maximum of three miscleavages were set using fixed carbamidomethyl modification, and the variable modifications oxidized methionine, protein N-terminal acetylation, and serine/threonine/tyrosine phosphorylation. A false discovery rate (FDR) of 1% was enforced across protein, peptide, and modification levels. Phosphorylated residues were localized using a site localization probability threshold of at least 0.75. Protein abundances were determined through label-free quantification with default settings. Database search outcomes underwent additional statistical analysis using Perseus (version 1.6.12). Statistical comparisons included two-tailed Student’s T-tests for comparing two means, one-way ANOVA followed by Dunnett’s test for comparing multiple groups to a single control, and two-way ANOVA followed by Dunnett’s post-hoc test for comparing two factors. Specific exceptions to these methods are detailed in the corresponding figure legends. Statistical analyses and plots were conducted using Perseus (version 1.6.12), Prism 10 (Graphpad Software Inc., version 10.0.0), or in house R scripts.

---

**Supplementary Figure S1.**
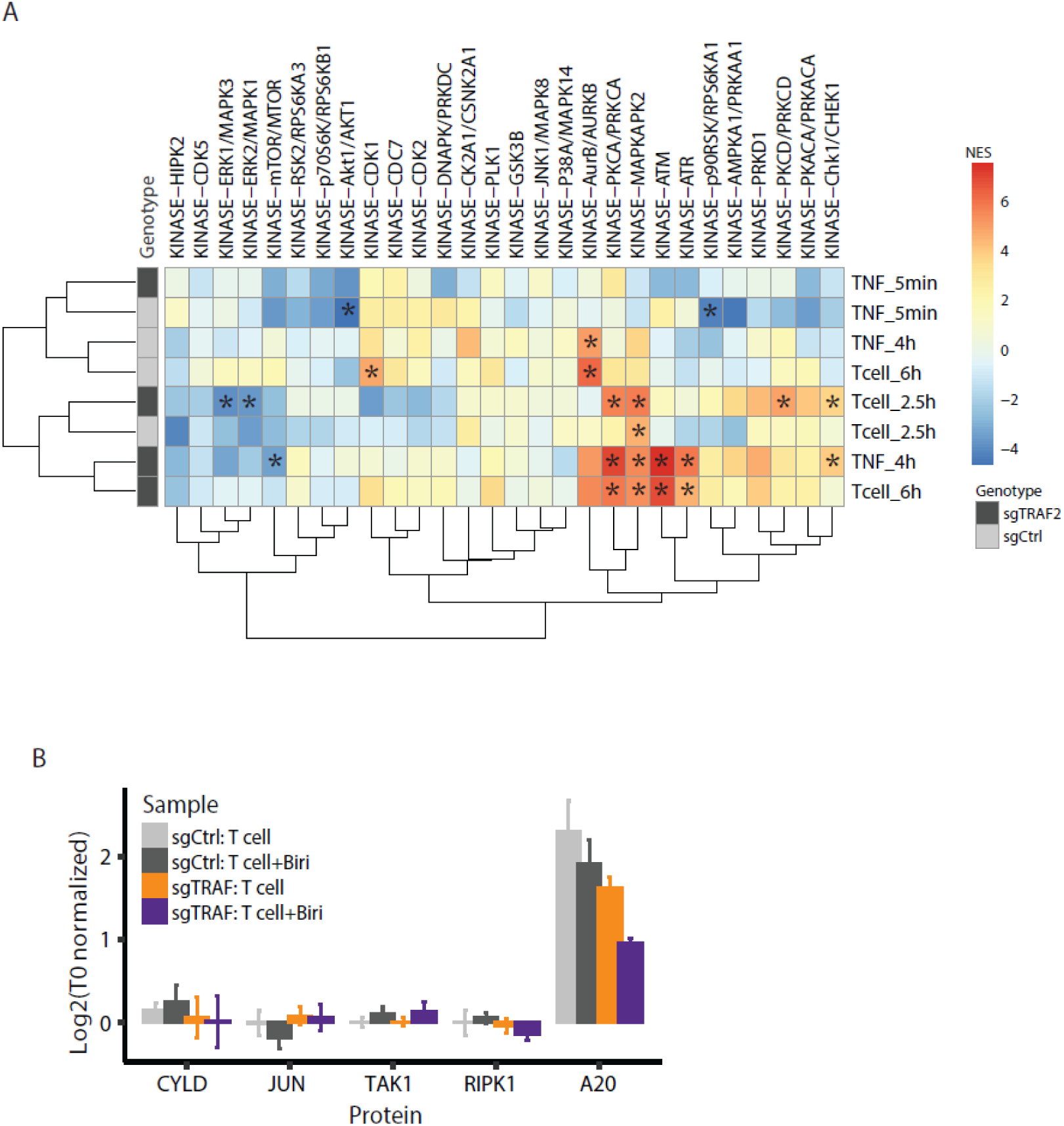
Kinase activation and abundance changes of select proteins in melanoma cells sensitized to T cell killing. **(A)** Hierarchal clustering of kinase activation signatures measured in sgCtrl or sgTRAF2 transduced D10 cells treated under the listed conditions. Sensitized sgTRAF2 cells show similar kinase activation to T cell and TNF treatment at later timepoints. Normalized Enrichment Scores (NES) calculated using PTM-SEA analysis [48]. Values represent average NES scores and * indicate significant NES with FDR < 5% in 3 biological replicates. **(B)** Protein abundance changes of dynamically regulated proteins at the phosphoproteome level following 6 hours of matched T cell co-culture using sgCtrl or sgTRAF2 transduced BLM cells with or without Birinapant treatment. Protein abundances are normalized to T0. Error bars represent standard deviation between biological replicates (n=3).

**Supplementary Figure S2.**
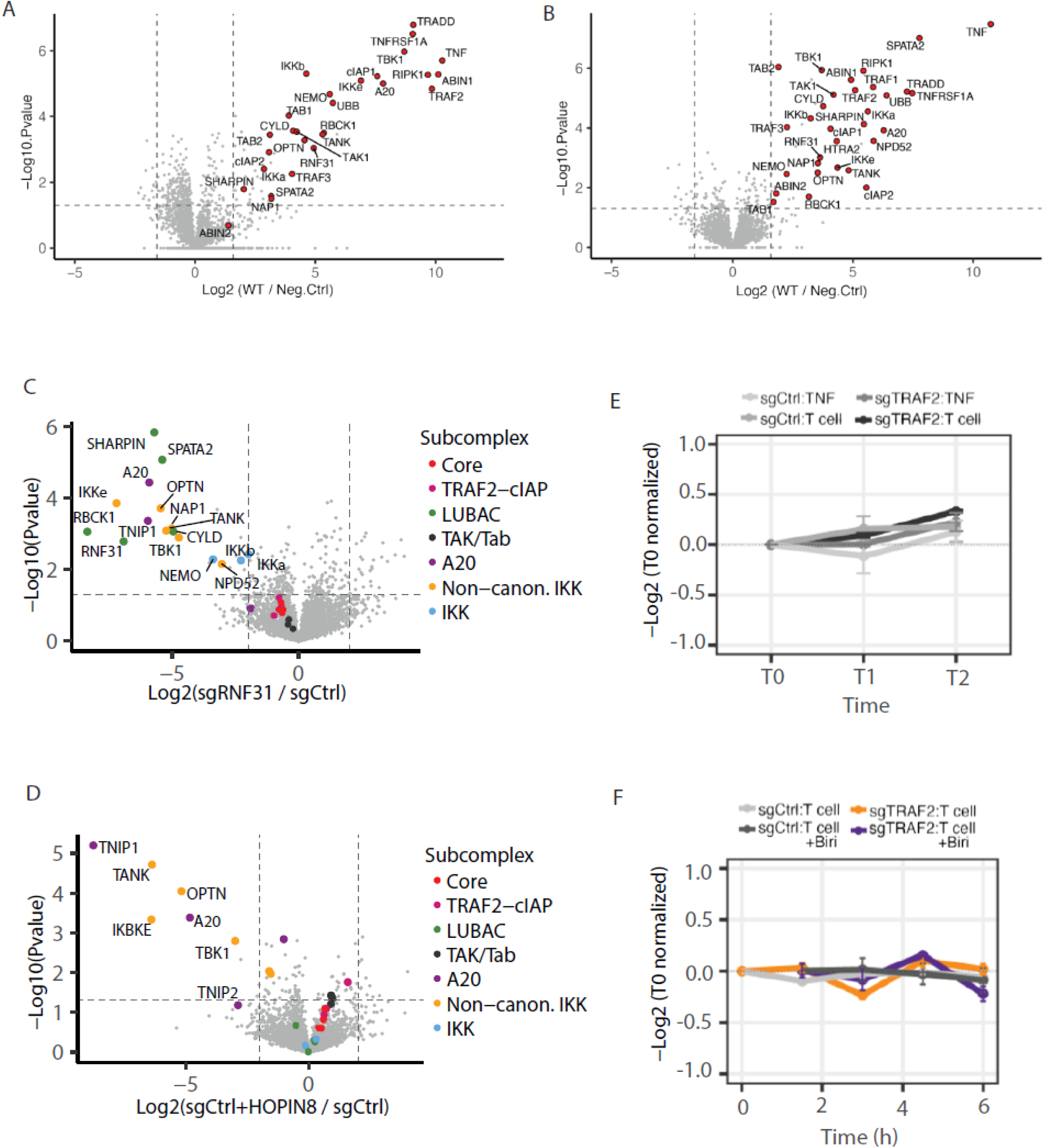
Enrichment of Complex I associated proteins using TNF AP-MS and RNF31 protein levels in melanoma treatment timecourse. **(A)** Relative abundance of Complex I associated proteins (red dots) in sgCtrl BLM cells treated with biotin-TNF (100ng/ml) vs. cells treated with untagged TNF (100ng/ml). Horizontal line indicates P-value < 0.05, vertical lines indicate fold-change > 3, n=3 biological replicates per condition. Protein abundance values were extracted from MQ output. **(B)** Same as (A) but data performed again in a second experimental set. **(C)** Relative abundance of Complex I proteins in sensitive vs resistant tumor cell. Tumor cells were sensitized to T cell-mediated killing by knocking out RNF31 and Complex I proteins were measured by TNF-biotin AP-MS. Complex I proteins are color coded by subcomplex. Horizontal line indicates P-value < 0.05, vertical lines indicate fold-change > 4, n=3 biological replicates per condition. **(D)** Same as in (D) but tumor cells were sensitized to T cell-mediated killing through treatment with RNF31 inhibitor HOPIN8. Figure S2C,2D were created with re-used data [42]. **(E)** RNF31 protein levels in sgCtrl or sgTRAF2 transduced D10 cells treated under the listed conditions. T1= 5min for TNF treatment and 2.5 h for T cell treatment. T2= 4 h for TNF treatment and 6h for T cell treatment. Error bars represent standard deviation between biological replicates (n=3). **(F)** RNF31 protein levels in sgCtrl or sgTRAF2 transduced BLM cells treated under the listed conditions. Error bars represent standard deviation between biological replicates (n=3).

**Supplementary Figure S3.**
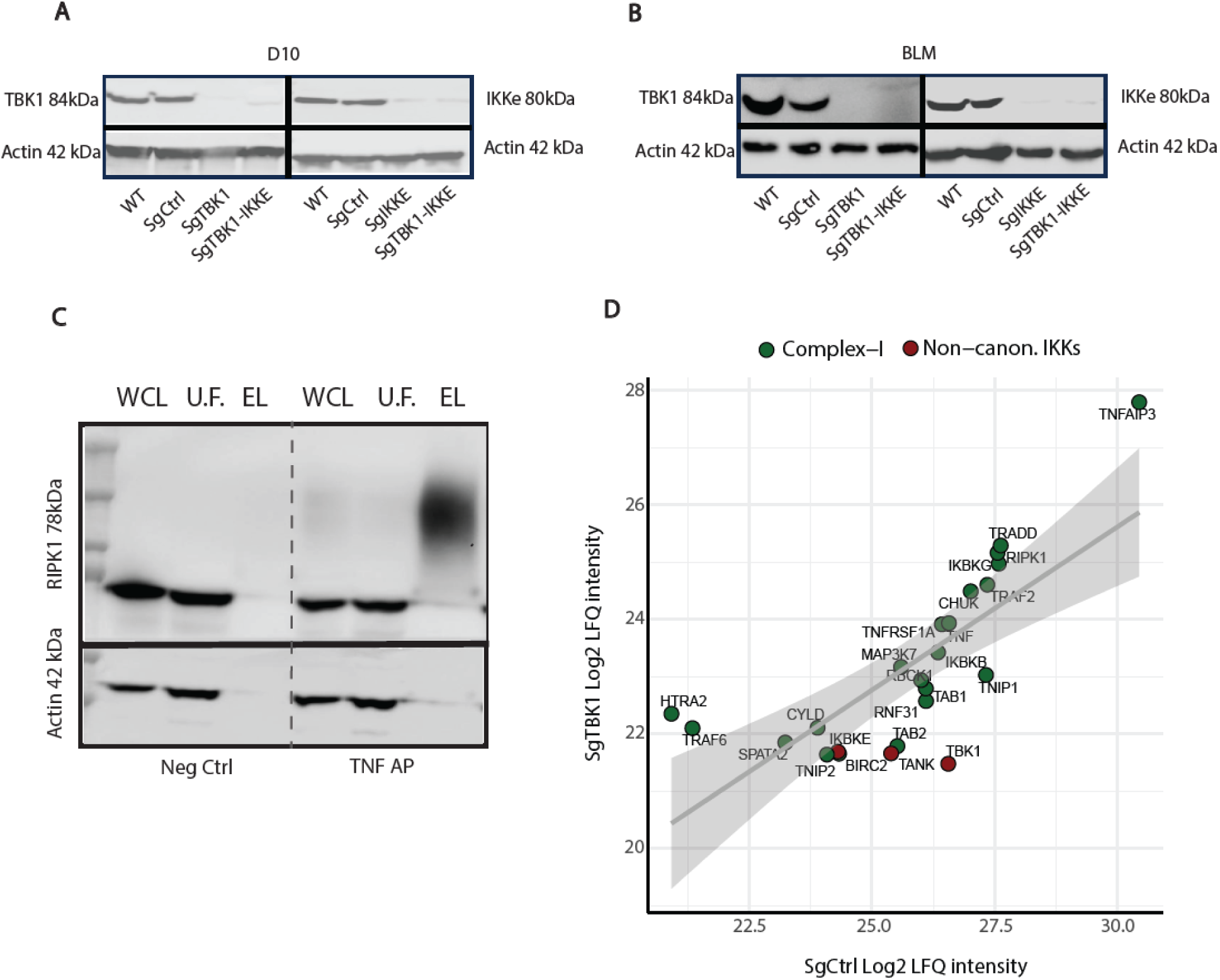
Validation of TNF-AP specificity and functional KOs of TBK1 and IKKE in two human melanoma cell lines. **(A,B)** Functional KOs of TBK1, IKKe in two human melanoma cell lines: BLM and D10. Tumor cells were harvested and whole cell lysates were blotted against TBK1 mAb and IKKe mAb. Β-actin was used as a loading control. **(C)** WB showing different steps of TNF-AP experiment in BLM cells. Cells were treated with TNF-biotin (TNF AP condition) or untagged TNF (Neg Ctrl condition) for 10min, followed by affinity purification as described in Fig2A. Fractions of each experimental step were blotted against RIPK1 mAb.. Β-actin was used as a loading control. WCL= whole cell lysate, U.F.= Unbound fraction, EL=elution using 5% SDS. (D) Correlation between relative protein abundance of Complex I proteins in SgTBK1 vs SgCtrl D10 cells. LFQ intensity values are Log2 transformed, mean normalized, and represent the average of 2 biological replicates per condition with missing values imputed.

**Supplementary Table S4.**
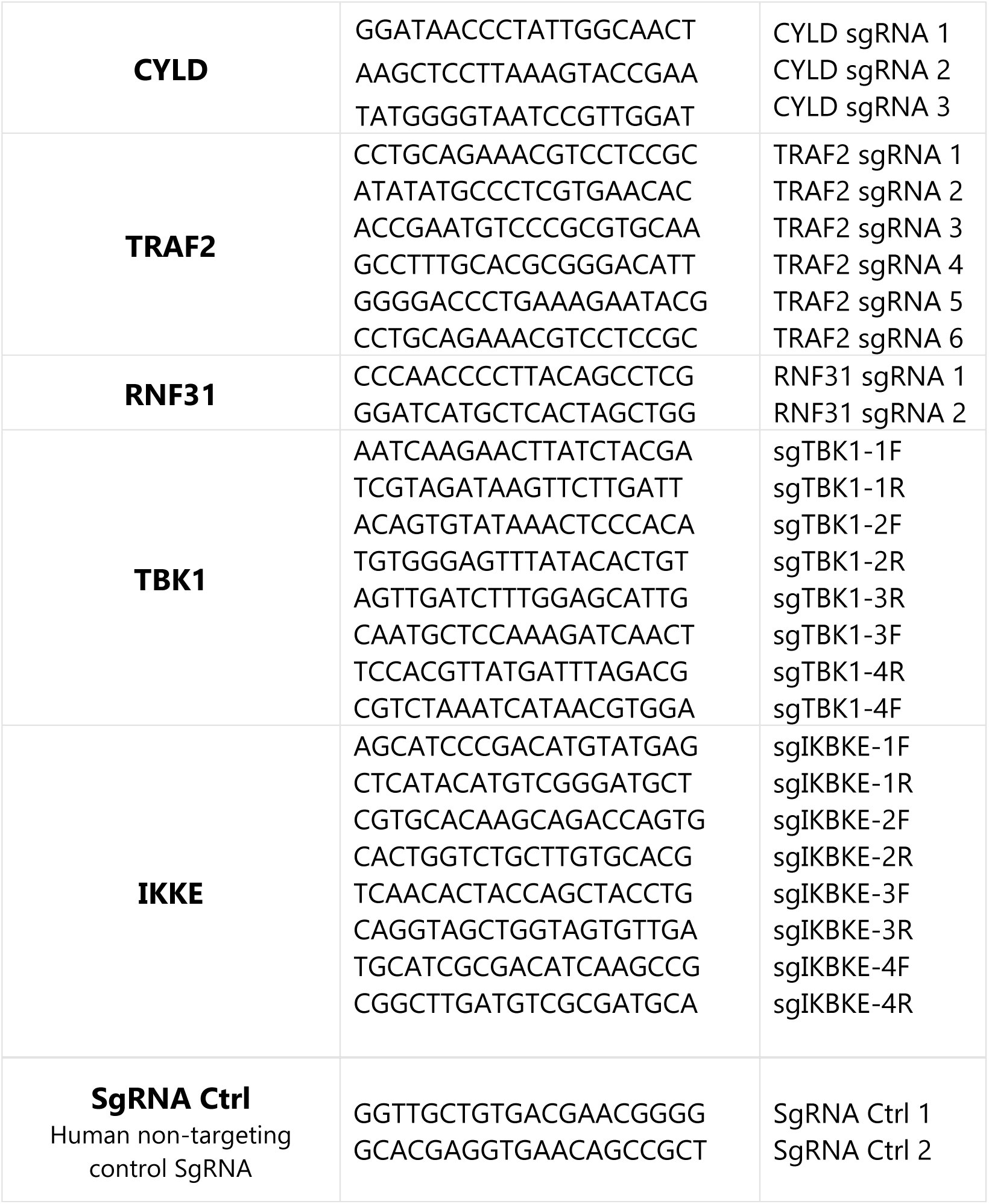
Target sgRNA oligonucleotide sequences CRISPR-mediated knockouts.

